# Pax6 maintains lens epithelial cell identity and coordinates secondary fiber cell differentiation

**DOI:** 10.1101/2025.10.23.684166

**Authors:** Barbora Antosova, Jana Smolikova, Anna Zitova, Jitka Lachova, Zbynek Kozmik

**Affiliations:** Laboratory of Transcriptional Regulation, Institute of Molecular Genetics, Academy of Sciences of the Czech Republic, Videnska 1083, Praha 4, 142 20, Czech Republic

**Author notes:** Corresponding author. Institute of Molecular Genetics, Academy of Sciences of the Czech Republic, Videnska 1083, Praha 4, 142 20, Czech Republic.

**Keywords:** Eye, Lens, Development, Pax6, Foxe3, Conditional knockout

## Abstract

Pax6 is a crucial regulator of vertebrate eye development, and its loss leads to the failure of lens placode formation. To investigate Pax6 function at successive stages of lens development, we employed the Cre-loxP system in combination with a novel *Foxe3-Cre* driver, which becomes active after the lens placode stage but prior to the onset of secondary fiber cell differentiation. The *Foxe3-Cre* enables efficient deletion of *Pax6* throughout the entire lens by embryonic day E12.5. Our study shows that *Pax6* loss causes a delay in lens differentiation, disrupts the lens epithelium, and produces a smaller lens that remains attached to the cornea, ultimately leading to a rudimentary lens in adulthood. Notably, Foxe3 persisted in the mutant lens epithelium despite Pax6 loss, while apoptosis and aberrant Sox2 upregulation occurred in the epithelium. Combined with the delayed onset of fiber cell differentiation, the abnormal anterior expansion of fiber cell differentiation regulators (c-Maf and Sox1), and the aberrant expression of cyclin D2, these results underscore the essential role of Pax6 in preserving lens epithelial identity and coordinating the transition to secondary fiber cell differentiation.

## Introduction

Genetic studies in multiple vertebrate models—human, mouse, rat, zebrafish, medaka, and *Xenopus*—have provided key insights into the role of Pax6 in visual system development (Glaser et al., 1994; Hill et al., 1991; Kleinjan et al., 2008; Matsuo et al., 1993; Mikula Mrstakova and Kozmik, 2024; Nakayama et al., 2015). Recent evidence suggests that the lens-specific functions of Pax6 are more deeply conserved across vertebrates than its roles in the retina (Mikula Mrstakova and Kozmik, 2024). The underlying reason for a conserved Pax6 requirement in vertebrate lens formation remains unresolved. This is particularly striking given the diversity of lens morphogenetic strategies among extant vertebrates—delamination in teleosts versus invagination in mammals and birds. In mammals, Pax6 is indispensable for Shroom-mediated epithelial bending and lens invagination (Plageman et al., 2010), a mechanism that is evidently unnecessary in fish, where the lens forms by delamination. These contrasts imply that, while the morphological route to a lens can vary, a core Pax6-dependent regulatory program may be preserved, with species-specific effectors executing distinct morphogenetic outcomes.

Lens development in mouse is a highly orchestrated complex proces starting with the thickening of head surface ectoderm and formation of lens placode. Subsequently, lens placode invaginates and form lens pit which separates from the surface ectoderm to become a lens vesicle. The anterior cells of the lens vesicle give rise to the lens epithelial cells, while the posterior cells elongate and produce primary lens fiber cells which fill the interior of the lens. The proliferation of lens epithelial cells produces new cells that upon differentiation in the equatorial region of the lens and cell cycle exit in transitional zone (TZ) generate secondary lens fiber cells, which contribute to lens growth throughout the life (Martinez and de Iongh, 2010; McAvoy et al., 1999). In this study, we use TZ for this functional band (cyclin D2/p27 high; onset of Sox1/Prox1/c-Maf) and reserve the term equatorial region for the broader anatomical belt (Cvekl and Ashery-Padan, 2014; Rowan et al., 2008; Shaham et al., 2009).

Although Pax6 null mice are anophthalmic, conditional alleles and stage-specific Cre drivers enabled dissection of tissue- and stage-specific roles during development (Ashery-Padan et al., 2000; Klimova and Kozmik, 2014; Marquardt et al., 2001; Oron-Karni et al., 2008; Raviv et al., 2014; Shaham et al., 2009; Sunny et al., 2022). In the mouse, Pax6 is expressed throughout all stages of lens development, except of terminally differentiated lens fiber cells (Shaham et al., 2012; Walther and Gruss, 1991). Lens placode formation intrinsically depends on Pax6: conditional deletion of *Pax6* at the placode stage using *Le-Cre* arrests lens development (Ashery-Padan et al., 2000). Similarly, disruption of the lens-specific shadow enhancers of *Pax6* phenocopies placodal Pax6 loss, blocking lens formation (Antosova et al., 2016). Pax6 expression persists in the anterior, undifferentiated lens epithelium and TZ, but when the equatorial epithelial cells proliferate, exit the cell cycle, and differentiate into secondary fiber cells, Pax6 expression is lost (Cvekl and Ashery-Padan, 2014; Duncan et al., 2004; Shaham et al., 2009). Conditional deletion of *Pax6* during secondary lens fiber cell differentiation using *MLR10-Cre* (Zhao et al., 2004) revealed a requirement of Pax6 for initiation of the lens fiber differentiation program (Shaham et al., 2009).

Foxe3 is expressed from the placode stage throughout the lens vesicle and then restricted to the anterior epithelium; it is required for epithelial proliferation and for vesicle detachment (Blixt et al., 2000; Brownell et al., 2000; Medina-Martinez et al., 2005). This expression pattern makes Foxe3 an attractive regulatory context for driving Cre in the post-placodal lens. In this study, we generated BAC transgenic mice expressing Cre recombinase under the control of the *Foxe3* gene’s cis-regulatory elements. We characterized both the spatial expression pattern of Cre and the efficiency of *Foxe3-Cre*– mediated recombination. Our findings indicate that *Foxe3-Cre* serves as a powerful tool for gene deletion throughout the lens following the lens placode stage. Using this novel driver, we employed a conditional mouse line carrying a floxed *Pax6* allele to investigate the effects of *Pax6* deletion from the lens vesicle stage onward and analyzed the resulting lens phenotype. This approach allowed us to interrogate Pax6 functions in epithelial maintenance and the transition to secondary fiber differentiation, prior to the time window targeted by *MLR10-Cre* (Shaham et al., 2009).

## Material and methods

### Ethics Statement

Experimental mice were housed and *in vivo* experiments were executed in compliance with the European Communities Council Directive of 24 November 1986 (86/609/EEC) and national and institutional guidelines. Experimental procedures for handling the mice were approved by the Animal Care Committee of the Institute of Molecular Genetics (AVCR 1979/2021 SOV II and AVCR 2599/2024 SOVII).

### Mouse strains

The following genetically modified mice were used in this study: ***Rosa26R*** (Soriano, 1999) (Jackson Laboratory, stock no. 003309), ***Pax6^f^*^l/fl^** (Klimova and Kozmik, 2014), and ***Foxe3-Cre*** transgenic mouse (this study, see below). A 152 kb Bacterial Artificial Chromosome (BAC) RP24-187N7 harboring the coding exon and the 5‘ and the 3‘ regions of the mouse *Foxe3* gene was purchased from Children’s Hospital Oakland Research Institute. To generate *Foxe3-Cre BAC*, the open reading frame of Cre::EGFP recombinase fusion was inserted into the exon containing the translation initiation codon of *Foxe3* using a method of BAC recombineering (Lee et al., 2001). Oligonucleotides used for BAC recombineering are listed in Supplementary Table 1. Modified *Foxe3-Cre* BAC DNA was isolated and used for pronuclear injection. Positive founder mice were identified using primers designed to recognize the CRE-EGFP junction. For the assesment of Cre activity, animals of the F2 and later generations derived from a single positive F0 founder were used for genetic crosses. Embryos exhibited reproducible Cre-mediated recombination, regardless of the breeding generation. The genotype of each mouse was determined by PCR analysis of tail DNA (oligonucleotides used are listed in Supplementary Table 1).

### X-Gal staining procedure

For the β-galactosidase assay, embryos were fixed in 2% PFA, followed by washing with a rinse buffer (0.1 M phosphate buffer pH 7.3, 2 mM MgCl2, 20 mM Tris pH 7.3, 0.01% sodium deoxycholate, and 0.02% Nonidet P-40) and incubated in X-Gal staining solution (rinse buffer supplemented with 5 mM potasium ferricyanide, 5 mM potassium ferrocyanide, 20 mM Tris pH 7.3, and 1 mg/ml X-gal) at 37°C for 2 hours and overnight at room temperature with shaking. For cryosections, embryos were re-fixed in 4% paraformaldehyde in 1% PBS, washed with PBS, cryopreserved in 30% sucrose, and embedded in OCT (Tissue Tek, Sakura Finetek). The frozen tissues were then sectioned at a thickness of 10-12µm.

### Histology and immunohistochemistry

Mouse embryos were staged by designation the noon of the day when the vaginal plug was observed as embryonic day 0.5 (E0.5). Embryos of the desired age were dissected, fixed in 4% paraformaldehyde (PFA) from 45 minutes up to overnight at 4°C, or fixed in 4% formaldehyde overnight at 4°C, washed 3 x 10 minutes in 1x PBS, and either cryopreserved in 30% sucrose, and frozen in OCT (Sakura), or embeded in paraffine for parafinne sections. The cryosections (10–12 μm) were permeabilized with PBT (PBS with 0.1% Tween), blocked with 10% BSA in PBT and incubated with the primary antibody (1% BSA in PBT) overnight at 4°C. Sections were then washed with PBS, incubated with a fluorescent secondary antibody (Life Technologies, 1:500) for one hour at room temperature, washed with PBS, counterstained with DAPI, and mounted in Mowiol. For histology analysis, paraffin sections (8µm) were deparaffinised, rehydrated with an ethanol series, and stained with Hematoxylim a Eosin. For immunofluorescence, paraffin sections were deparaffinised, rehydrated with an ethanol series, followed by antigen retrieval, performed by heat-treating the sections in sodium citrate buffer (10mM sodium citrate buffer, 0.05% Tween 20, pH 6.0). The sections were then blocked in 10% BSA in PBT for 30 minutes, followed by overnight incubation with the primary antibody in 1% BSA in PBT at 4°C. The following day, sections were washed 3x for 10 minutes in 1x PBS, incubated with the secondary antibody in 1% BSA in PBT for one hour at room temperature, washed again 3x for 10 minutes in PBT, counterstained with DAPI (1µg/ml), and mounted in Mowiol (Sigma). The images were acquired using a Leica SP5 confocal microscope, or an Andor Dragonfly 503 – spinning disk confocal microscope. The following primary antibodies were used: anti-GFP (Life Technologies, A11122), anti-Pax6 (Covance, PRB-278P), anti-Foxe3 (Peter Carlsson), anti-cCaspase3 (Cell Signaling, #9664), anti-cyclin D2 (Santa-Cruz, sc-452), anti-Sox1 (Santa Cruz, sc-17317), anti-c-Maf (Bethyl Laboratories, BL662), anti-Sox2 (Santa Cruz, sc-17320).

## Results

### Generation of *Foxe3-Cre* transgenic mice and characterization of Cre activity

For the generation of *Foxe3-Cre* transgenic mice, we selected a 152kb BAC clone (# RP24-187N7) covering the *Foxe3* locus likely containing cis-regulatory sequences ensuring proper spatio-temporal expression. Employing the method of BAC recombineering (Lee et al., 2001), a cassette carrying fusion of the Cre recombinase and EGFP was introduced into the first ATG of the *Foxe3* gene (Fig. 1A). The modified BAC clone was then used for pronuclear injections, and founders were screened for the presence of the BAC. A single transgenic line was established by breeding with *C57Bl/6* mice. To confirm Cre activity, native EGFP protein was detected in the lens in *Foxe3-Cre* embryos at E11.5 (Fig. 1B), coinciding with the stage when Foxe3 is expressed in entire lens vesicle (Blixt et al., 2000). Real-time transgene expression was monitored using anti-GFP antibody on sections of the developing lens in entire lens vesicle at E12.0 (Fig. 1C) and in lens epithelial cells at E13.5 (Fig. 1D). To visualize Cre recombinase activity in cellular descendants of cells in which *Cre* expression occurs at any time during development, *Foxe3-Cre* transgenic mice was mated with *ROSA26R* reporter strain (Soriano, 1999). X-gal staining of lacZ (β-galactosidase) in *Foxe3-Cre; ROSA26R* embryos confirmed Cre activity in the lens pit and also in nasolacrimal groove and in the most caudal dorsolateral parts of the diencephalon at E10.5 (Fig. 1F), whereas, at E9.5, Cre is not yet expressed (Fig. 1E). At E12.5, robust *Foxe-Cre* activity was observed in the lens and diencephalon, with weaker staining in the future conjunctival epithelium (Fig. 1G). X-gal stained cryosections of the eye region of E10.5 and E12.5 *Foxe3-Cre; ROSA26R* embryos confirmed strong and specific *Foxe3-Cre* activity in the lens pit, later resulting in lens epithelium and lens fiber cells, as well as in the future conjunctival epithelium (Fig. 1H, I). The spatio-temporal expression pattern of *Foxe3-Cre* confirms its suitability for specific gene deletion in the lens.

**Figure 1.**
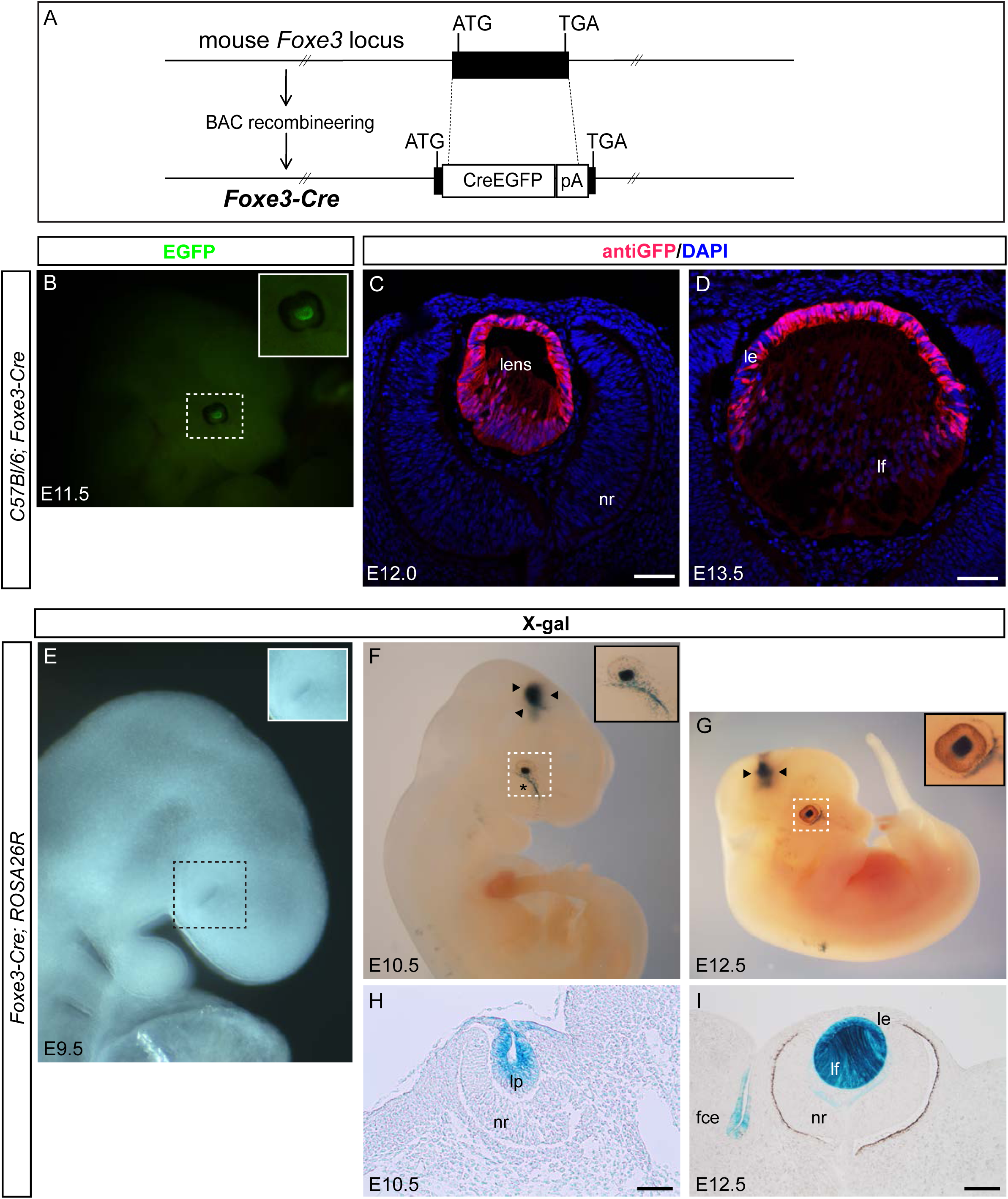
Generation of *Foxe3-Cre* BAC and characterisation of Cre expression pattern. (**A**) Schematic representation illustrating the modifiacation of a BAC clone containing the mouse *Foxe3* gene by BAC recombineering. A cassette containing the coding sequence of Cre recombinase fused to the coding sequence of EGFP was inserted into the first ATG of the *Foxe3* gene. Black boxes indicate exons. **(B)** Real-time monitoring of *Cre* expression using native EGFP in an E11.5 *Foxe3-Cre* whole-mount embryo. The inset provides a higher magnification of the eye region (boxed), showing EGFP+ lens vesicle. **(C-D)** Sections through the eye region co-stained with DAPI (blue) showing real-time *Cre* expression by immunofluorescent detection of EGFP in the entire lens at E12.0 and later in lens epithelial cells at E13.5. **(E-I)** Assessment of *Foxe3-Cre* activity using the *ROSA26R* reporter mouse. **(E-G)** Whole-mounts or **(H,I)** cryosections stained with X-gal to detect lacZ at indicated stages, demonstrating Cre activity. **(E)** At E9.5, Cre is not yet expressed in *Foxe3-Cre; ROSA26R* embryos. **(F)** At E10.5, Cre transgene expression was observed in the lens vesicle, nasolacrimal groove (asterisk), and diencephalon (arrowheads). **(G)** At E12.5, strong Cre expression was detected in the lens and persisted in the diencephalon. The insets provide higher magnification of the eye region (boxed). **(H, I)** Cryosections of the eye region in *Foxe3-Cre; ROSA26R* embryos show *Cre* activation specifically in the lens vesicle at E10.5 and in the entire lens at E12.5, when it was also detected in the future conjuctival epithelium. lens; nr, neural retina; le, lens epithelium; lf, lens fibers; lp, lens pit; fce, future conjunctival epithelium Scale bars: (C, D) 50 µm, (G) 50 µm, (H) 100 µm.

### *Foxe3-Cre* efficiently deleted Pax6 in the lens

To assess the effectiveness and specificity of the novel *Foxe3-Cre* in conditional gene deletion, we generated *Foxe3-Cre; Pax6^fl/fl^* transgenic mice. This transgene enabled us to study the role of Pax6 in lens development, particulary between the lens vesicle stage (using *Le-Cre* (Ashery-Padan et al., 2000)) and E14.5 (using *MLR10-Cre* (Shaham et al., 2009)). To determine the initiation of Pax6 deletion in *Foxe3-Cre; Pax6^fllfl^* transgenic lens, we monitored Pax6 loss by immunostaining at varios stages (Fig. 2A-F). Initially, we focused on stage E11.5, when *Foxe3-Cre* had been active at least one day. At E11.5 Pax6 was detected throughout the entire lens vesicle in controls (Fig. 2A), while in *Foxe3-Cre; Pax6^fllfl^* embryos, the Pax6 immunostaining began to weaken in the posterior lens vesicle (Fig. 2B). By E12.5, Pax6 was entirely absent from the *Foxe3-Cre; Pax6^fl/fl^* lenses (Fig. 2D, F), aligning phenotypically with smaller lens size in mutant embryos (Fig. 2D, F).

**Figure 2.**
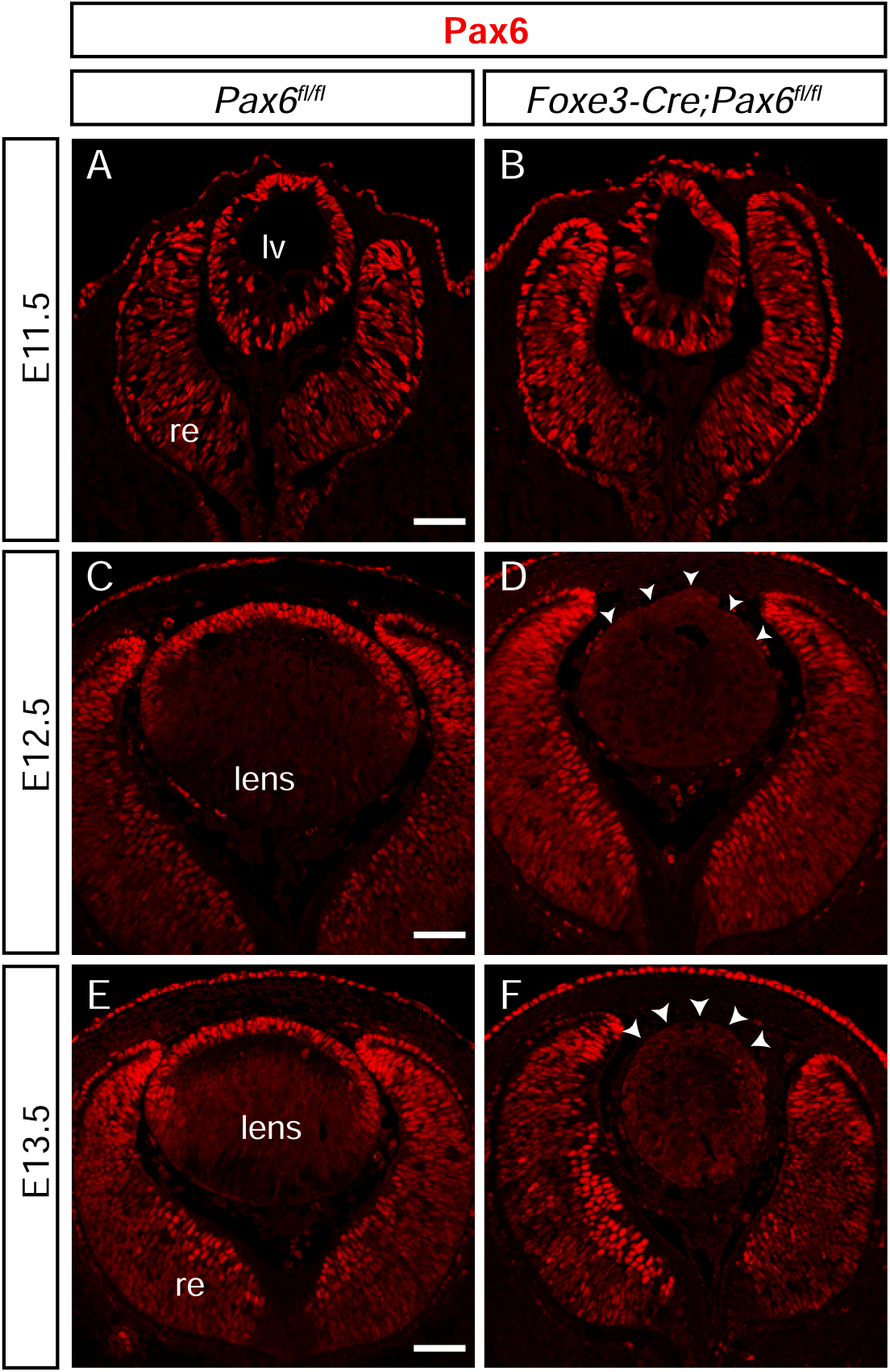
*Foxe3-Cre*-mediated Pax6 inactivation in the developing lens. **(A-F)** Immunofluorescent detection of Pax6 at indicated stages of lens development in control (*Pax6^fl/fl^*) and *Foxe3-Cre; Pax6^fl/fl^* cryo-(E11.5) or paraffin sections (E12.5, E13.5). **(A, B)** During the lens vesicle stage, Pax6 levels start to decrease in the lens vesicle of *Foxe3-Cre; Pax6^fl/fl^* transgenic embryos **(B)** compared to controls **(A)**. **(D, F)** In E12.5 and E13.5 *Foxe3-Cre; Pax6^fl/fl^* lenses, Pax6 is not detected in the entire lens, while **(C, E)** it is present in lens epithelial cells in controls. lv, lens vesicle; re, retina. Scale bar: (A-F) 50 µm.

### Deletion of *Pax6* using *Foxe3-Cre* resulted in a smaller lens that remained connected to the cornea

Adult *Foxe3-Cre; Pax6^fl/fl^*mice expressed extreme microphthalmia (Fig. 3B) compared to their *Pax6^fl/fl^* littermates (Fig. 3A). To characterize the morphological defects of the *Foxe3-Cre; Pax6^fl/fl^*lenses, we performed histological analysis on eyes from 4-week-old adults (Fig. 3C, D) and embryos at various developmental stages (Fig. 3E-L). This analysis revealed a rudimentary lens in 4-week-old mutant mice (Fig. 3D). The lens of *Foxe3-Cre; Pax6^fl/fl^*was vacuolated, with the anterior lens epithelium failing to separate from the cornea, and a lens stalk was present (Fig. 3D’, D’’). Additionally, the mutant exhibited a thinner corneal epithelium (Fig. 3D’, D’’) and abnormal retinal lamination (Fig. 3D). To uncover gradual development of this terminal lens phenotype, we focused on embryonic stages of lens development. At E11.5, when *Foxe3-Cre* was already active but Pax6 was still present in the lens vesicle (Fig. 2B), we did not observe significant morphological defects in *Foxe3-Cre; Pax6^fl/^*^fl^ lenses (Fig. 3F). Later at E12.5, when Pax6 was completely lost from the lens vesicle in *Foxe3-Cre; Pax6^fl/fl^* mutants (Fig. 2D), we observed a reduction in lens size in *Foxe3-Cre; Pax6^fl/fl^* embryos (Fig. 3H), which progressed at E14.5 and E16.5 (Fig. 3J, L). Furthermore, at E12.5, *Foxe3-Cre; Pax6^fl/fl^*exhibited an obvious persistent lens vesicle, suggesting a delay in lens development (Fig. 3H). At E14.5 (Fig. 3J), similarly as at E16.5 (Fig. 3L), we also observed remaining fiber cell nuclei in the lens posterior in *Foxe3-Cre; Pax6^fl/fl^* lenses, along with vacuoles in the anterior and a persistent connection of the lens to the cornea (Fig. 3J, L). At E16.5, abnormal eye morphology and reduced eye size were clearly visible in *Foxe3-Cre; Pax6^fl/fl^* whole-mount embryos compared to littermate controls (Fig. 3M, N).

**Figure 3.**
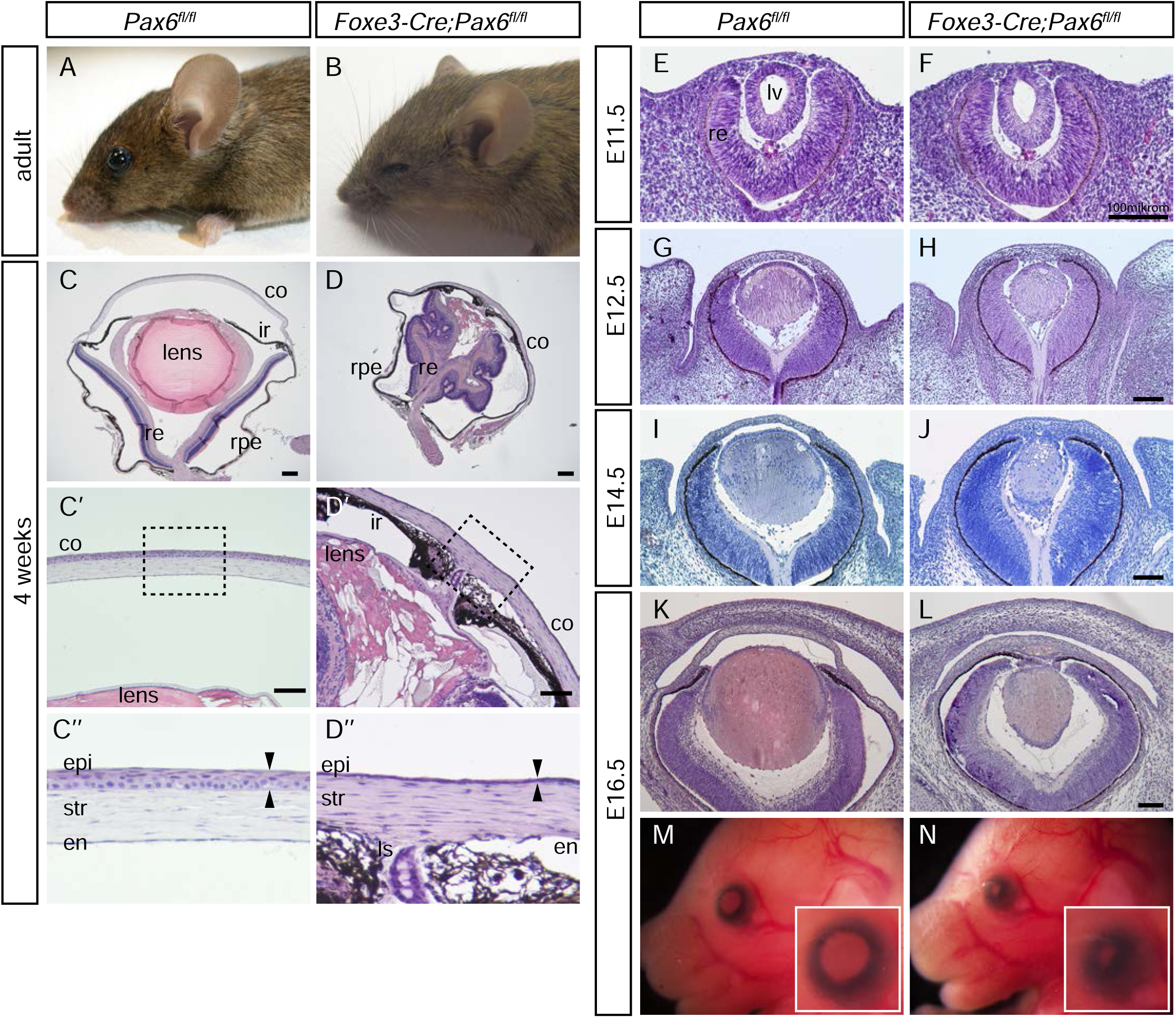
Morphological consequences of *Pax6* deletion in lens vesicle stage. **(A,B)** Adult *Foxe3-Cre; Pax6^fl/fl^* mice exhibit extreme microphthalmia compared to controls (*Pax6^fl/fl^*). **(C-J)** Histological analysis of eye or eye region on sections of control and *Foxe3-Cre; Pax6^fl/fl^* mice or embryos. **(D)** The eyes of 4-week-old *Foxe3-Cre; Pax6^fl/fl^*mice are smaller compared to controls **(C)**, featuring a rudimentary vacuolated lens with a persisting connection to the cornea, which is attached to the iris. **(D)** The retina of *Foxe3-Cre; Pax6^fl/fl^*is folded and retains some lamination. **(D’, D’’)** The cornea of 4-week-old *Foxe3-Cre; Pax6^fl/fl^*is affected, attached to the iris, and the corneal epithelium is thinner (arrowheads) compared to controls **(C’, C’’)**. **(D’’)** A detailed view of the boxed region of cornea shows in *Foxe3-Cre; Pax6^fl/fl^* a thinner corneal epithelium, abnormal corneal stroma, and corneal endothelium attached to the iris, contrasting with the normal corneal morphology in control **(C’’)**. **(F)** At E11.5, the lens vesicle of *Foxe3-Cre; Pax6^fl/fl^* is comparable to the control **(E)**, although it stays in contact with the surface ectoderm. **(H)** From E12.5, the lens of *Foxe3-Cre; Pax6^fl/fl^* is smaller than control **(G)**, and this reduction in lens size progresses at E14.5 **(J)**, and E16.5 **(L)**. **(J)** At E14.5, vacuoles are already present in the *Foxe3-Cre; Pax6^fl/fl^* lens, and nuclei of lens fiber cells persist in the lens posterior. **(M, N)** At E16.5 abnormal eye morphology and reduced eye size were clearly visible in *Foxe3-Cre; Pax6^fl/fl^* whole-mount embryos compared to controls. co, cornea; ir, iris; re, retina; rpe, retina pigmented epithelium; epi, corneal epithelium; str, corneal stroma; en, corneal endothelium; ls, lens stalk; lv, lens vesicle. Scale bars: (C, D) 200 µm, (C’, D’, E-H) 100 µm.

### Foxe3 persisted in the lens epithelium of *Foxe3-Cre; Pax6^fl/fl^* embryonic lenses

Previous studies have shown the sensitivity of Foxe3 to the presence of Pax6 in the lens (Antosova et al., 2016; Blixt et al., 2007). Therefore, we examined Foxe3 expression in *Foxe3-Cre; Pax6^fl/fl^* mutants. At E11.5, when Pax6 was still present in the lens vesicle, there was no obvious change in Foxe3 expression in *Foxe3-Cre; Pax6^fl/fl^*lens (Fig. 4B) compared to the control (Fig. 4A). Surprisingly, at E12.5 and E13.5, when Pax6 was already absent in the lens, Foxe3 was not lost in the lens epithelium of *Foxe3-Cre; Pax6^fl/fl^* mutants either (Fig. 4D, F). However, at 12.5 Foxe3 was not present in all the lens epithelial cells of *Foxe3-Cre; Pax6^fl/fl^* lens (Fig. 4D) as it was in the control (Fig. 4C). One day later at E13.5, considering the already evident aberrant morphology of the epithelium, the reduction in Foxe3 in *Foxe3-Cre; Pax6^fl/fl^* lenses was not as pronounced (Fig. 4F). These findings indicate that the eye phenotype observed in *Foxe3-Cre; Pax6^fl/fl^* mice is not a result of Foxe3 loss, but rather reflects a specific role of Pax6—or possibly another key transcription factor distinct from Foxe3—in lens development. In addition, the data suggest that Pax6 is necessary only for the initial activation of *Foxe3*, whereas beyond the lens vesicle stage, Foxe3 expression can be maintained independently of Pax6.

**Figure 4.**
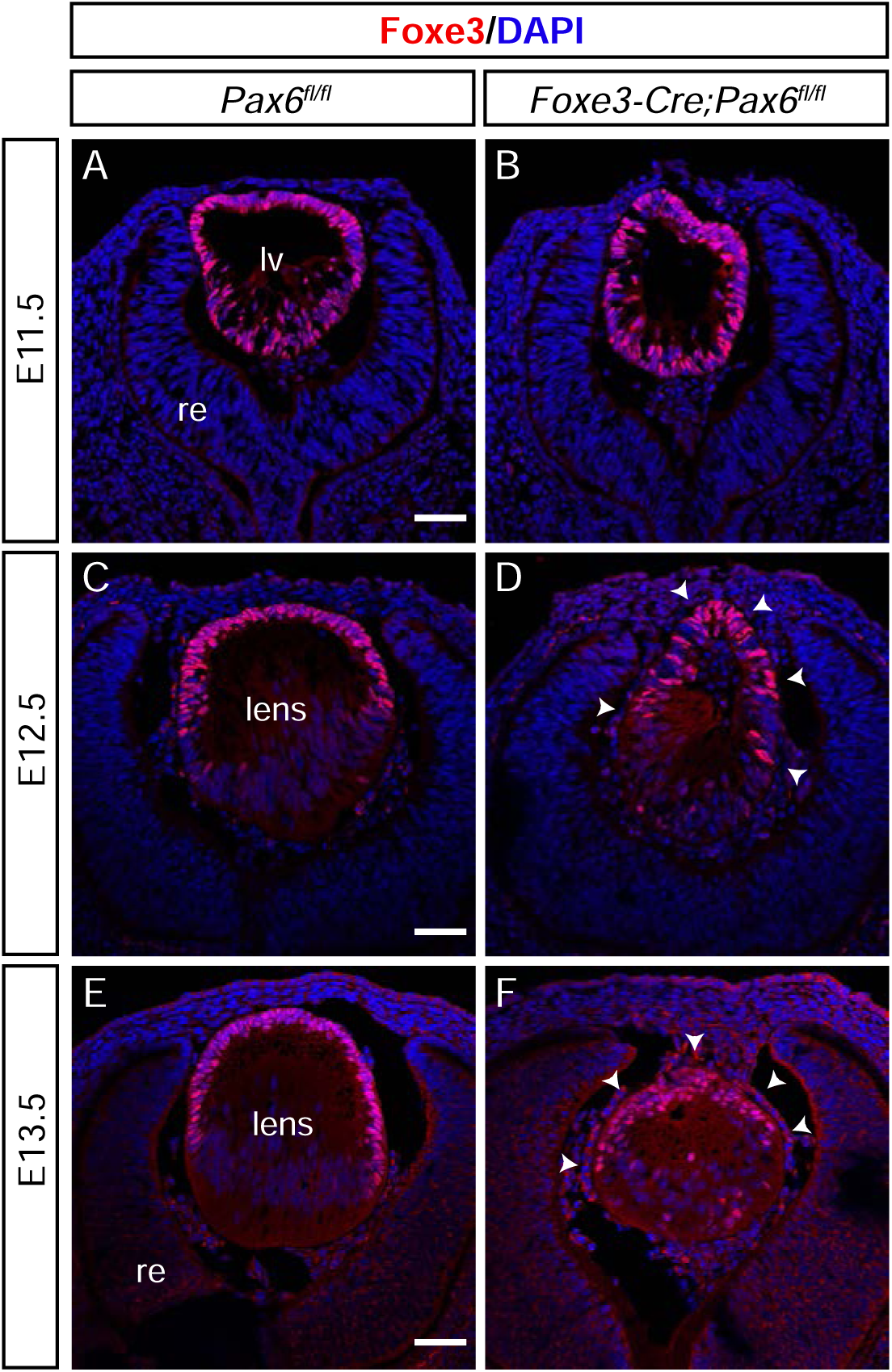
Pax6 is not essential for the maintenance of Foxe3 expression in the lens. **(A-F)** Immunofluorescent detection of Foxe3 counterstained with DAPI on cryosections. **(A, B)** At E11.5, Foxe3 is expressed in the entire lens vesicle in wildtype and in *Foxe3-Cre; Pax6^fl/fl^* lenses. **(C, E)** At E12.5 and E13.5, Foxe3 expression is restricted to the lens epithelium in controls. **(D, F)** Despite altered lens morphology, Foxe3 is expressed in the lens anterior of *Foxe3-Cre; Pax6^fl/f^* mutants (arrowheads). lv, lens vesicle; re, retina. Scale bar: (A-F) 50 µm.

### The lens epithelium and fiber cell differentiation were affected in *Foxe3-Cre; Pax6^fl/fl^* lenses

The first obvious manifestation of Pax6 deletion in *Foxe3-Cre; Pax6^fl/fl^* eyes was a reduction in overall lens size and a delay in lens development (Fig. 3H), as evidenced by the persistent lens vesicle observed at E12.5 and E13.5 (Fig. 3H, J). To investigate whether the smaller lens size in *Foxe3-Cre; Pax6^fl/fl^* mutants was partially due to apoptosis, we performed immuno-fluorescent detection of cleaved caspase 3 (cCas3) (Fig. 5A, B). At E13.5, apoptotic cells were detected in the prospective lens epithelium of *Foxe3-Cre; Pax6^fl/fl^* lenses (Fig. 5B). Furthermore, remaining fiber cell nuclei were observed posterior to the lens equator even at E14.5 of *Foxe3-Cre; Pax6^fl/fl^*lenses (Fig. 3J). This suggested, apart from delayed development, aberrant fiber cell differentiation in the equatorial region, consistent with a previously described defect in fiber cell differentiation following Pax6 loss in later developing lens (Shaham et al., 2009). Consequently, we investigated the expression of cyclin D2, which is specifically expressed in the transitional zone in controls at E13.5 (Fig. 5C) – the equatorial region of lens where fiber cells differentiate, and the expression pattern of c-Maf a Sox1 (Fig. 5E-H), essential regulators of fiber cell differentiation (Kim et al., 1999; Nishiguchi et al., 1998). In *Foxe3-Cre; Pax6^fl/fl^* lenses at E13.5, we detected cyclin D2 positive nuclei localized in the lens posterior (Fig. 5D), corresponding more to the expression pattern of cyclin D2 at E11.5 in lens vesicle (Rowan et al., 2008). Additionally, c-Maf-positive cell nuclei were observed in the lens posterior of E13.5 *Foxe3-Cre; Pax6^fl/fl^* lenses (Fig. 5F), aligning more with the c-Maf expression pattern at E12.5 lens (Kerr et al., 2012), when primary fiber cell differentiation occurrs (Cvekl and Ashery-Padan, 2014). In controls at E13.5 (Fig. 5C, E), secondary fiber cells have been already formed (Cvekl and Ashery-Padan, 2014). This suggests that lens development was delayed due to the affected differentiation of secondary fiber cells in *Foxe3-Cre; Pax6^fl/fl^* mutants. Finally, we analyzed the expression of Sox1, which, in E13.5 controls, was expressed exclusively in fiber cells (Fig. 5G), whereas in *Foxe3-Cre; Pax6^fl/fl^* mutants, its expression was detected anteriorly from the equatorial region in lens epithelial cells (Fig. 5H).

**Figure 5.**
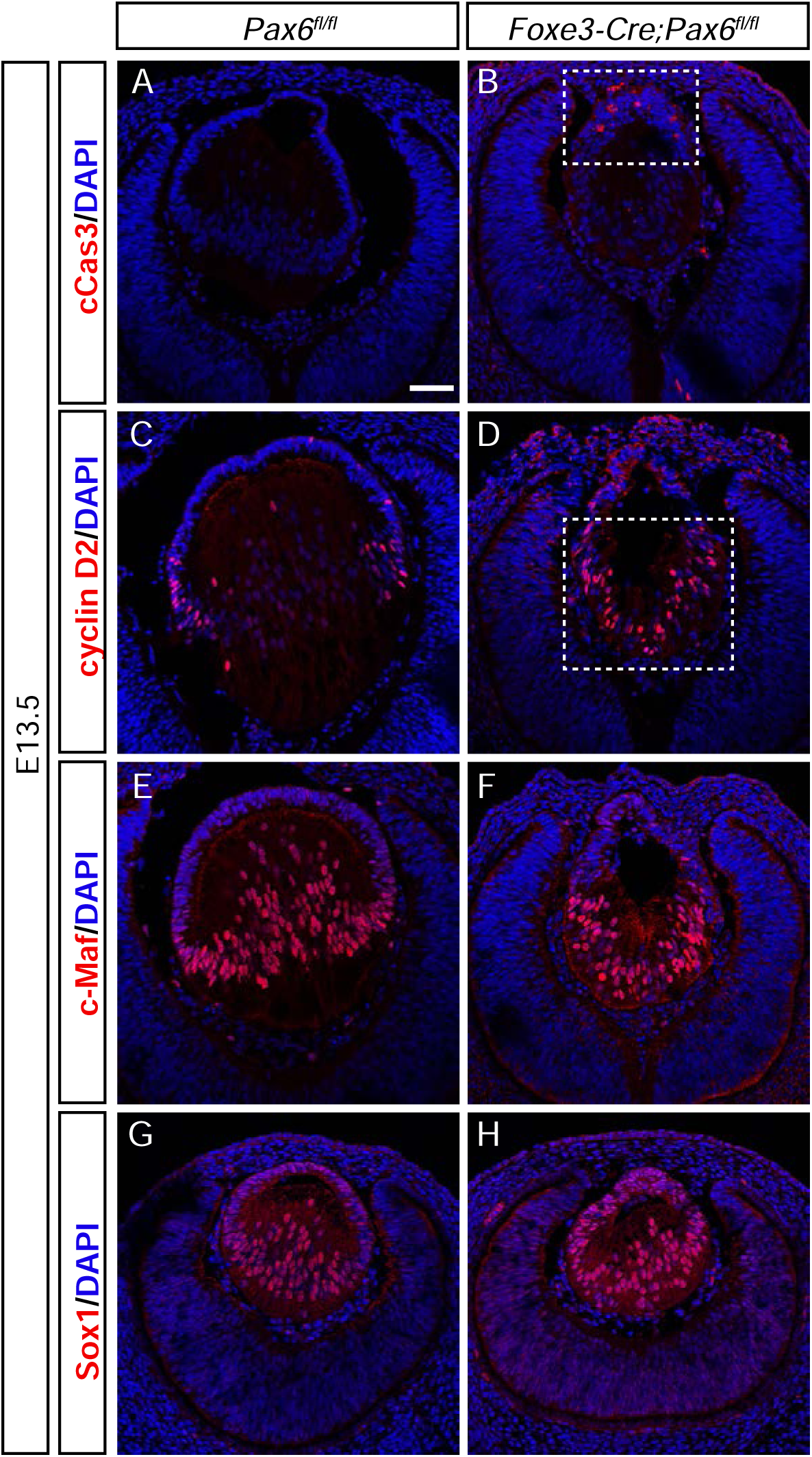
The lens epithelium is affected in *Foxe3-Cre; Pax6^fl/fl^* lenses. **(A-H)** Immunofluorescent detection of apoptosis **(A, B)**, and fiber cell differentiation markers **(C-H)** counterstained with DAPI in control (*Pax6^fl/fl^*) and *Foxe3-Cre; Pax6^fl/fl^* lenses at the indicated developmental stages. **(B)** Apoptotic cells are marked with cCasp3 in *Foxe3-Cre; Pax6^fl/fl^* lens epithelium. **(D)** Cyclin D2 positive cells are detected in the posterior of the lens vesicle of *Foxe3-Cre; Pax6^fl/fl^* mutants (also delayed morphological development of the lens is detected), whereas it is present only in the transitional zone of E13.5 control lenses **(C)**. **(F)** The fiber cell differentiation marker c-Maf is detected in the lens posterior of *Foxe3-Cre; Pax6^fl/fl^* lenses, while **(H)** Sox1 positive cells expanse to the lens anterior from the equatorial region in *Foxe3-Cre; Pax6^fl/fl^* mutants. re, retina. Scale bar: (A-H) 50 µm.

Previously, Pax6 has been shown to bind enhancer sequences of *Sox2* and activate Sox2 expression in lens cells (Inoue et al., 2007; Lengler et al., 2005). Sox2 is expressed in lens placode, and lens vesicle until E12.5, when it is apparently downregulated in lens (Kamachi et al., 1998; Nishiguchi et al., 1998). Notably, *Pax6* ablation using *MLR10-Cre* resulted in an increase in Sox2 expression in the lens transitional zone (Shaham et al., 2009), though not in the anterior lens epithelium. Therefore, we investigated Sox2 expression following Pax6 loss. In *Foxe3-Cre; Pax6^fl/fl^* lenses, we observed increase of Sox2 expression in the anterior lens at both E12.5 and at E13.5 (Fig. 6B, D, F), in contrast to controls (Fig. 6A, C, E), where Sox2 was typically reduced in the lens at this stage. Furthermore, Pax6/Sox2 double-staining nicely confirmed the mutually exclusive expression of Pax6 and Sox2 in the lens epithelium/lens anterior (Fig. 6E, F).

**Figure 6.**
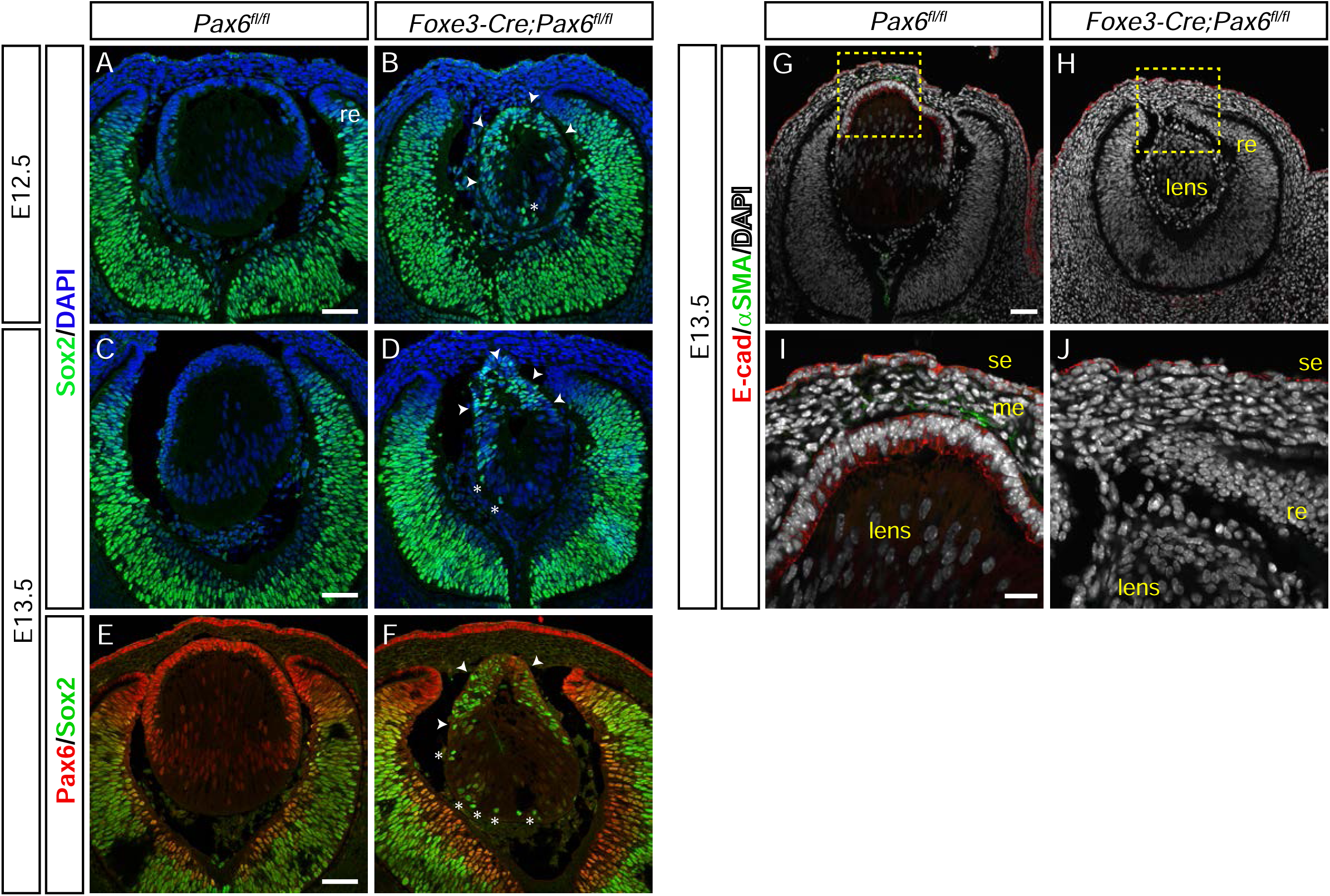
The lens epithelium of *Foxe3-Cre; Pax6^fl/fl^* lenses loses its epithelial character. **(A-J)** Immunoflurescent detection of Sox2 **(A-D)**, Pax6 and Sox2 double-staining **(E, F)**, and E-cadherin and α-SMA **(G-J),** counterstained with DAPI, in controls (Pax6*^fl/fl^*) and *Foxe3-Cre; Pax6^fl/fl^* lenses at the indicated developmental stages. **(A-F)** In E12.5 **(B)** and E13.5 **(D)**, *Foxe3-Cre; Pax6^fl/fl^* lenses show strong upregulation of Sox2 in the lens epithelium (arrowheads) where Pax6 is lost **(F)**, and ectopic expression in the lens posterior (asterisks) **(B, D, F)**. **(G-J)** E-cadherin is detected in lens epithelium of control lenses **(G, I)**, and in surface ectoderm of both control and *Foxe3-Cre; Pax6^fl/fl^* embryos **(G-J)**, but its expression is lost from the anterior lens of *Foxe3-Cre; Pax6^fl/fl^* mutants **(H, J)**. **(I, J)** α-SMA is detected in the mesenchyme of controls **(I)**, but not in the lens anterior of *Foxe3-Cre; Pax6^fl/fl^* embryos **(J)**. me, mesenchyme; se, surface ectoderm; re, retina. Scale bars: (A-H) 50 µm, (I, J) 20 µm.

To further determine whether *Foxe3-Cre; Pax6^fl/fl^* lens epithelium retained or lost its epithelial character, we analyzed the expression of E-cadherin, an epithelial marker expressed in the surface ectoderm, and later in lens vesicle and lens epithelium (Pontoriero et al., 2009), and α-smooth muscle actin (α-SMA), a key marker of epithelial-to-mesenchymal transition (EMT), at embryonic day 13.5 (E13.5), (Fig. 6G-J). E-cadherin expression was lost in the lens anterior of *Foxe3-Cre; Pax6^fllfl^* mutants (Fig. 6H, J), while it remained present in the overlying surface ectoderm of both control and *Foxe3-Cre; Pax6^fl/fl^* embryos (Fig. 6 I, J). At E13.5, no α-SMA signal was detected in the lens anterior of *Foxe3-Cre; Pax6^fl/fl^* (Fig. 6J), indicate that EMT was not occuring in this development stage.

Taken together, fiber cell differentiation in *Foxe3-Cre; Pax6^fl/fl^* lenses was delayed while lens epithelial cells lost their epithelial character. These findings suggest a crucial role for Pax6 in the proper specification of lens epithelial cells and the differentiation of secondary fiber cells.

## Discussion and conslusions

### *Foxe3-Cre* as a tool for lens-vesicle specific gene ablation

Here, we generated a *Foxe3-Cre* transgenic line that enables efficient deletion of floxed alleles throughout the lens vesicle and epithelium, beginning after lens placode formation and prior to the onset of secondary fiber cell differentiation. Notably, this developmental window has not been specifically or cleanly targeted by previously available Cre lines such as *Le-Cre*, *LR-Cre*, *P0-3.9GFPCre*, or *MLR10-Cre* (Ashery-Padan et al., 2000; Kreslova et al., 2007; Lam et al., 2019; Rowan et al., 2008; Zhao et al., 2004). For example, *Le-Cre* (*Pax6 P0/EEL* based) deletes at the placode and arrests lens development; confounding CRE toxicity exists in homozygotes (Ashery-Padan et al., 2000; Lam et al., 2019). *LR-Cre* and *P0-3.9GFPCre* target placode/early vesicle but may have broader surface ectoderm or even neuroretina activity (Kreslova et al., 2007; Lam et al., 2019; Rowan et al., 2008). Finally, *MLR10-Cre* deletes later, during secondary fiber differentiation (Shaham et al., 2009; Zhao et al., 2004).

The *Foxe3-Cre* line opens a practical “second-window” for genetic analysis—after placode formation yet before secondary fiber differentiation—that cleanly decouples lens induction from epithelial maturation and the TZ handoff, inviting systematic tests of regulators that are indispensable earlier and thus inaccessible with placodal Cre lines. For example, prior work points to signaling pathways or chromatin modulators that gate *Pax6* output (Cvekl and Zhang, 2017). *Foxe3-Cre* now enables these pathways to be interrogated in the same developmental frame in which Pax6 maintains epithelial character, thereby resolving hierarchy—which processes are upstream of, parallel to, or downstream from Pax6. This should also clarify whether adhesion failure and lens-stalk persistence are primary defects or secondary consequences of transcriptional mispatterning, and why penetrance varies with timing and dosage. The approach has direct translational relevance, bridging mouse phenotypes with *PAX6*- and *FOXE3*-linked anterior segment disease, including aniridia and Peters’ anomaly. Ultimately, this window should define the minimal regulatory module required for lens epithelial integrity and lens–cornea separation, while revealing points of genetic compensation that explain variability across strains and contexts.

### Pax6 role in lens epithelial identity and secondary fiber cell differentiation

As previously described by Cvekl and Ashery-Padan (Cvekl and Ashery-Padan, 2014), primary lens fiber cell differentiation begins around embryonic day 12.5 (E12.5), while by E13.5–E14.5, primary fiber cell elongation is largely completed and the formation of secondary lens fiber cells is initiated. In our study, the lens phenotype observed following *Pax6* deletion coincides precisely with this developmental transition. The differentiation of lens fiber cells appears to stall shortly after E12.5, suggesting that the loss of *Pax6* prevents the proper switch from primary to secondary fiber cell differentiation. Morphologically, this developmental arrest is evident as a failure of the lens fibers to elongate normally and organize into the layered structure typical of later stages. These findings indicate that Pax6 plays a continuous and indispensable role in orchestrating the progression of fiber cell differentiation beyond the primary fiber cell stage. In parallel, the lens epithelium exhibits pronounced molecular changes upon *Pax6* loss, most notably the reactivation of *Sox2*. Sox2 is a well-known pluripotency-associated transcription factor that, during normal lens development, is expressed in early stages but becomes progressively downregulated in the lens epithelium by approximately E12.5 (Arnold et al., 2011). Its re-expression in the *Pax6*-deficient epithelium is striking, as it suggests a partial reversion of epithelial cells toward a less differentiated, more progenitor-like state. This aberrant Sox2 upregulation likely reflects a loss of epithelial identity and highlights a regulatory role of Pax6 in maintaining the differentiation status of the lens epithelium.

Compared with the later *MLR10-Cre; Pax6^fl/fl^* model—where *Pax6* loss yields a smaller lens primarily from defective fiber differentiation and Sox2 upregulation in the TZ (Shaham et al., 2009) — our *Foxe3-Cre; Pax6^fl/fl^* mutants exhibit an earlier block: the TZ fails to form, Sox2 does not rise in the TZ but instead surges anteriorly, and differentiation stalls prior to stable secondary fiber onset. This timing shift explains the distinct phenotypes and underscores that the developmental stage and domain of Pax6 inactivation determine outcome. In our model, variable lens-stalk persistence and reduced lens size align with Pax6 dosage sensitivity during vesicle detachment (Sey/+; aniridia) and the need for cadherin-mediated adhesion (Blixt et al., 2007; Blixt et al., 2000; Pontoriero et al., 2009).

Mechanistically, we observe anterior apoptosis, loss of E-cadherin, a Sox2 increase in epithelium, anterior expansion of c-Maf/Sox1, and a cyclin D2 shift from the equator toward posterior/vesicle-like domains. These changes indicate that Pax6 restrains a progenitor-like Sox2 state, stabilizes epithelial junctions, and is required to establish the TZ where secondary fibers normally initiate (Rowan et al., 2008; Shaham et al., 2009). Consistent with prior work, Sox1/c-Maf can persist without Pax6, yet are insufficient to drive fiber differentiation on their own—mirroring our earlier-window mispatterning (Shaham et al., 2009). Finally, *Foxe3-Cre* deletion across the lens vesicle/early epithelium (∼E12.5) reveals that Pax6 is dispensable for short-term Foxe3 maintenance, but it is essential for epithelial identity and the timely transition to secondary fiber differentiation, with incomplete stalk penetrance likely reflecting the timimg of deletion relative to vesicle closure.

In summary, we propose a model in which Pax6 acts as a bimodal regulator in this window: (i) sustaining epithelial identity (E-cadherin, Sox2 modulation, survival), and (ii) priming the TZ to proceed toward secondary fiber differentiation (setting cyclin D2 domain, allowing c-Maf/Sox1 to regionalize). Loss of Pax6 destabilizes the epithelium (apoptosis, adhesion loss), collapses proper TZ architecture, and locks the lens into a delayed, primary-fiber–like trajectory that culminates in rudimentary, vacuolated remnants adherent to the cornea.

## Acknowledgements

This work was supported by the institutional funding (RVO68378050-KAV-NPUI) and grant by the Czech Science Foundation GA24-12482S (to Z.K.). We acknowledge the Light Microscopy Core Facility, IMG, Prague, Czech Republic, supported by MEYS – LM2023050, MEYS – CZ.02.1.01/0.0/0.0/18_046/0016045 and MEYS – CZ.02.01.01/00/23_015/0008205, for their support with the confocal imaging presented herein.

## Author contributions

BA Writing – original draft, Writing – review & editing, Conceptualization, Investigation, Methodology, Data curation.

JS Writing – review & editing, Investigation, Methodology.

AZ Writing – review & editing, Investigation, Methodology.

ZK: Writing – review & editing, Writing – original draft, Conceptualization, Investigation, Methodology, Resources, Data curation, Funding acquisition, Project administration.

## Declaration of generative AI and AI-assisted technologies in the manuscript preparation process

During the preparation of this work the authors used ChatGPT in order to improve English phrasing. After using this tool, the authors reviewed and edited the content as needed and take full responsibility for the content of the published article.

---

**Supplementary Table S1.**
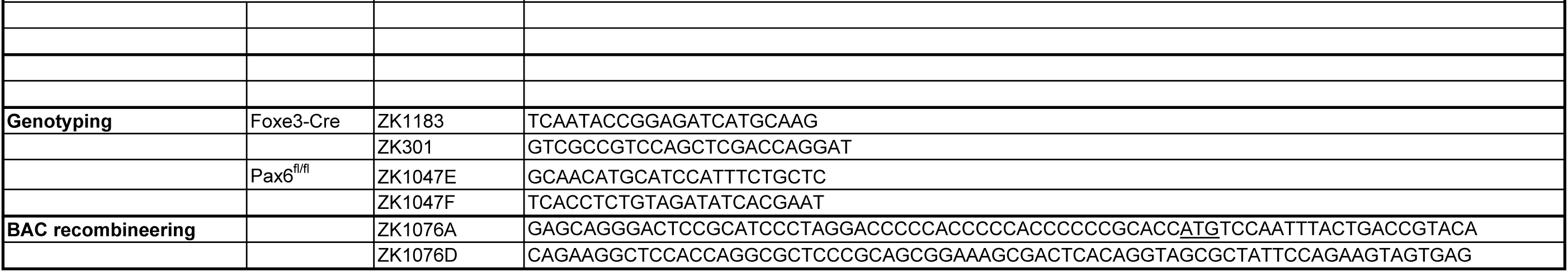
Oligonucleotides.

## References

Antosova, B., Smolikova, J., Klimova, L., Lachova, J., Bendova, M., Kozmikova, I., Machon, O., Kozmik, Z., 2016. The Gene Regulatory Network of Lens Induction Is Wired through Meis-Dependent Shadow Enhancers of Pax6. PLoS Genet 12, e1006441.

Arnold, K., Sarkar, A., Yram, M.A., Polo, J.M., Bronson, R., Sengupta, S., Seandel, M., Geijsen, N., Hochedlinger, K., 2011. Sox2(+) adult stem and progenitor cells are important for tissue regeneration and survival of mice. Cell Stem Cell 9, 317–329.

Ashery-Padan, R., Marquardt, T., Zhou, X., Gruss, P., 2000. Pax6 activity in the lens primordium is required for lens formation and for correct placement of a single retina in the eye. Genes Dev 14, 2701–2711.

Blixt, A., Landgren, H., Johansson, B.R., Carlsson, P., 2007. Foxe3 is required for morphogenesis and differentiation of the anterior segment of the eye and is sensitive to Pax6 gene dosage. Developmental biology 302, 218–229.

Blixt, A., Mahlapuu, M., Aitola, M., Pelto-Huikko, M., Enerback, S., Carlsson, P., 2000. A forkhead gene, FoxE3, is essential for lens epithelial proliferation and closure of the lens vesicle. Genes Dev 14, 245–254.

Brownell, I., Dirksen, M., Jamrich, M., 2000. Forkhead Foxe3 maps to the dysgenetic lens locus and is critical in lens development and differentiation. Genesis 27, 81–93.

Cvekl, A., Ashery-Padan, R., 2014. The cellular and molecular mechanisms of vertebrate lens development. Development 141, 4432–4447.

Cvekl, A., Zhang, X., 2017. Signaling and Gene Regulatory Networks in Mammalian Lens Development. Trends Genet 33, 677–702.

Duncan, M.K., Xie, L., David, L.L., Robinson, M.L., Taube, J.R., Cui, W., Reneker, L.W., 2004. Ectopic Pax6 expression disturbs lens fiber cell differentiation. Invest Ophthalmol Vis Sci 45, 3589–3598.

Glaser, T., Jepeal, L., Edwards, J.G., Young, S.R., Favor, J., Maas, R.L., 1994. PAX6 gene dosage effect in a family with congenital cataracts, aniridia, anophthalmia and central nervous system defects. Nat Genet 7, 463–471.

Hill, R.E., Favor, J., Hogan, B.L., Ton, C.C., Saunders, G.F., Hanson, I.M., Prosser, J., Jordan, T., Hastie, N.D., van Heyningen, V., 1991. Mouse small eye results from mutations in a paired-like homeobox-containing gene. Nature 354, 522–525.

Inoue, M., Kamachi, Y., Matsunami, H., Imada, K., Uchikawa, M., Kondoh, H., 2007. PAX6 and SOX2-dependent regulation of the Sox2 enhancer N-3 involved in embryonic visual system development. Genes Cells 12, 1049–1061.

Kamachi, Y., Uchikawa, M., Collignon, J., Lovell-Badge, R., Kondoh, H., 1998. Involvement of Sox1, 2 and 3 in the early and subsequent molecular events of lens induction. Development 125, 2521–2532.

Kerr, C.L., Huang, J., Williams, T., West-Mays, J.A., 2012. Activation of the hedgehog signaling pathway in the developing lens stimulates ectopic FoxE3 expression and disruption in fiber cell differentiation. Invest Ophthalmol Vis Sci 53, 3316–3330.

Kim, J.I., Li, T., Ho, I.C., Grusby, M.J., Glimcher, L.H., 1999. Requirement for the c-Maf transcription factor in crystallin gene regulation and lens development. Proc Natl Acad Sci U S A 96, 3781–3785.

Kleinjan, D.A., Bancewicz, R.M., Gautier, P., Dahm, R., Schonthaler, H.B., Damante, G., Seawright, A., Hever, A.M., Yeyati, P.L., van Heyningen, V., Coutinho, P., 2008. Subfunctionalization of duplicated zebrafish pax6 genes by cis-regulatory divergence. PLoS Genet 4, e29.

Klimova, L., Kozmik, Z., 2014. Stage-dependent requirement of neuroretinal Pax6 for lens and retina development. Development 141, 1292–1302.

Kreslova, J., Machon, O., Ruzickova, J., Lachova, J., Wawrousek, E.F., Kemler, R., Krauss, S., Piatigorsky, J., Kozmik, Z., 2007. Abnormal lens morphogenesis and ectopic lens formation in the absence of beta-catenin function. Genesis 45, 157–168.

Lam, P.T., Padula, S.L., Hoang, T.V., Poth, J.E., Liu, L., Liang, C., LeFever, A.S., Wallace, L.M., Ashery-Padan, R., Riggs, P.K., Shields, J.E., Shaham, O., Rowan, S., Brown, N.L., Glaser, T., Robinson, M.L., 2019. Considerations for the use of Cre recombinase for conditional gene deletion in the mouse lens. Hum Genomics 13, 10.

Lee, E.C., Yu, D., Martinez de Velasco, J., Tessarollo, L., Swing, D.A., Court, D.L., Jenkins, N.A., Copeland, N.G., 2001. A highly efficient Escherichia coli-based chromosome engineering system adapted for recombinogenic targeting and subcloning of BAC DNA. Genomics 73, 56–65.

Lengler, J., Bittner, T., Munster, D., Gawad Ael, D., Graw, J., 2005. Agonistic and antagonistic action of AP2, Msx2, Pax6, Prox1 AND Six3 in the regulation of Sox2 expression. Ophthalmic Res 37, 301–309.

Marquardt, T., Ashery-Padan, R., Andrejewski, N., Scardigli, R., Guillemot, F., Gruss, P., 2001. Pax6 is required for the multipotent state of retinal progenitor cells. Cell 105, 43–55.

Martinez, G., de Iongh, R.U., 2010. The lens epithelium in ocular health and disease. Int J Biochem Cell Biol 42, 1945–1963.

Matsuo, T., Osumi-Yamashita, N., Noji, S., Ohuchi, H., Koyama, E., Myokai, F., Matsuo, N., Taniguchi, S., Doi, H., Iseki, S., et al., 1993. A mutation in the Pax-6 gene in rat small eye is associated with impaired migration of midbrain crest cells. Nat Genet 3, 299–304.

McAvoy, J.W., Chamberlain, C.G., de Iongh, R.U., Hales, A.M., Lovicu, F.J., 1999. Lens development. Eye (Lond) 13 (Pt 3b), 425–437.

Medina-Martinez, O., Brownell, I., Amaya-Manzanares, F., Hu, Q., Behringer, R.R., Jamrich, M., 2005. Severe defects in proliferation and differentiation of lens cells in Foxe3 null mice. Mol Cell Biol 25, 8854–8863.

Mikula Mrstakova, S., Kozmik, Z., 2024. Genetic analysis of medaka fish illuminates conserved and divergent roles of Pax6 in vertebrate eye development. Front Cell Dev Biol 12, 1448773.

Nakayama, T., Fisher, M., Nakajima, K., Odeleye, A.O., Zimmerman, K.B., Fish, M.B., Yaoita, Y., Chojnowski, J.L., Lauderdale, J.D., Netland, P.A., Grainger, R.M., 2015. Xenopus pax6 mutants affect eye development and other organ systems, and have phenotypic similarities to human aniridia patients. Dev Biol 408, 328–344.

Nishiguchi, S., Wood, H., Kondoh, H., Lovell-Badge, R., Episkopou, V., 1998. Sox1 directly regulates the gamma-crystallin genes and is essential for lens development in mice. Genes Dev 12, 776–781.

Oron-Karni, V., Farhy, C., Elgart, M., Marquardt, T., Remizova, L., Yaron, O., Xie, Q., Cvekl, A., Ashery-Padan, R., 2008. Dual requirement for Pax6 in retinal progenitor cells. Development 135, 4037–4047.

Plageman, T.F., Jr., Chung, M.I., Lou, M., Smith, A.N., Hildebrand, J.D., Wallingford, J.B., Lang, R.A., 2010. Pax6-dependent Shroom3 expression regulates apical constriction during lens placode invagination. Development 137, 405–415.

Pontoriero, G.F., Smith, A.N., Miller, L.A., Radice, G.L., West-Mays, J.A., Lang, R.A., 2009. Co-operative roles for E-cadherin and N-cadherin during lens vesicle separation and lens epithelial cell survival. Dev Biol 326, 403–417.

Raviv, S., Bharti, K., Rencus-Lazar, S., Cohen-Tayar, Y., Schyr, R., Evantal, N., Meshorer, E., Zilberberg, A., Idelson, M., Reubinoff, B., Grebe, R., Rosin-Arbesfeld, R., Lauderdale, J., Lutty, G., Arnheiter, H., Ashery-Padan, R., 2014. PAX6 regulates melanogenesis in the retinal pigmented epithelium through feed-forward regulatory interactions with MITF. PLoS Genet 10, e1004360.

Rowan, S., Conley, K.W., Le, T.T., Donner, A.L., Maas, R.L., Brown, N.L., 2008. Notch signaling regulates growth and differentiation in the mammalian lens. Dev Biol 321, 111–122.

Shaham, O., Menuchin, Y., Farhy, C., Ashery-Padan, R., 2012. Pax6: a multi-level regulator of ocular development. Prog Retin Eye Res 31, 351–376.

Shaham, O., Smith, A.N., Robinson, M.L., Taketo, M.M., Lang, R.A., Ashery-Padan, R., 2009. Pax6 is essential for lens fiber cell differentiation. Development 136, 2567–2578.

Soriano, P., 1999. Generalized lacZ expression with the ROSA26 Cre reporter strain. Nature genetics 21, 70–71.

Sunny, S.S., Lachova, J., Dupacova, N., Kozmik, Z., 2022. Multiple roles of Pax6 in postnatal cornea development. Dev Biol 491, 1–12.

Walther, C., Gruss, P., 1991. Pax-6, a murine paired box gene, is expressed in the developing CNS. Development 113, 1435–1449.

Zhao, H., Yang, Y., Rizo, C.M., Overbeek, P.A., Robinson, M.L., 2004. Insertion of a Pax6 consensus binding site into the alphaA-crystallin promoter acts as a lens epithelial cell enhancer in transgenic mice. Invest Ophthalmol Vis Sci 45, 1930–1939.

